# BIN1 overexpression rescues cardiac but not skeletal muscle defects in a mouse model of caveolinopathy

**DOI:** 10.1101/2025.09.26.678914

**Authors:** Alice Moncheaux, Alain Guimond, Coralie Spiegelhalter, Nadia Messaddeq, Chadia Nahy, Jocelyn Laporte

**Affiliations:** Institute of Genetics and Molecular and Cellular Biology (IGBMC), INSERM U1258, CNRS UMR7104, University of Strasbourg, Illkirch, France; Institut Clinique de la Souris, PHENOMIN-ICS, Illkirch, France

**Author notes:** correspondence to Jocelyn Laporte: IGBMC, 1 rue Laurent Fries, 67404 Illkirch, France. **Authors contribution:** JL conceived and supervised the project. AG performed cardiac function measurements. CS, NM and CN performed electron microscopy. AM conducted and analyzed all other experimental data, including *in vivo* motor assessments, force measurements, histology and molecular investigations. AM and JL wrote the manuscript. **Disclosure:** JL patented BIN1 expression for the treatment of centronuclear myopathies.

**Keywords:** Cavin, amphiphysin, ryanodine receptor, myopathy, cardiomyopathy, long QT syndrome, rippling muscle disease, centronuclear myopathy, T-tubule, gene therapy

## Abstract

**Background:** Mutations in *CAV3*, encoding caveolin-3, cause caveolinopathies, rare genetic disorders affecting both skeletal and cardiac muscle. Caveolin-3 contributes to T-tubule formation and excitation-contraction coupling. BIN1 (amphiphysin 2), a membrane-shaping protein critical for T-tubule integrity, has shown therapeutic promise in congenital myopathies and heart dysfunction. To date, there are no therapy for caveolinopathies.

**Methods:** We evaluated the therapeutic impact of BIN1 overexpression in *Cav-3* knockout mice, a model recapitulating key features of human caveolinopathy. We assessed skeletal and cardiac function, T-tubule morphology, mitochondria, and gene expression using histological, physiological, and molecular approaches.

**Results:** We found *Cav-3*^*-/-*^ mice displayed skeletal muscle weakness, T-tubule disorganization, and mitochondrial abnormalities, alongside cardiac diastolic dysfunction and myofibrillar disarray. While BIN1 overexpression failed to improve skeletal muscle strength, T-tubule structure, or fiber atrophy, it corrected nuclear positioning and partially restored mitochondrial markers. In contrast, BIN1 robustly rescued cardiac performance, restoring end-diastolic volume, cardiac output, and sarcomeric integrity. Expression profiling revealed greater dysregulation of excitation-contraction coupling and atrogene pathways in skeletal than in cardiac muscle. Cavin-4, a BIN1-interacting protein, was selectively dysregulated in *Cav3-/-* muscle, suggesting a mechanistic barrier to BIN1-mediated rescue in this tissue.

**Conclusions:** These findings identify tissue-specific differences in the molecular consequences of caveolin-3 loss and demonstrate that BIN1 overexpression effectively rescues cardiac, but not skeletal, manifestations of caveolinopathy. Our results support BIN1 as a promising gene therapy target for inherited cardiomyopathies, while highlighting the need for alternative strategies in skeletal muscle.

## INTRODUCTION

Loss-of-function mutations in the gene *CAV3*, encoding for caveolin-3, are responsible for heterogeneous and diverse muscular disorders gathered under the term caveolinopathies^1^. This group encompasses both skeletal muscle diseases, such as distal myopathy (MIM# 614321) and rippling muscle disease (MIM#606072), as well as cardiac conditions, including hypertrophic cardiomyopathy (MIM#192600)^2^ and long QT syndrome^3^ (MIM#611818). Most prominent clinical signs include muscle rippling, exercise intolerance, calf hypertrophy and elevated CK levels in the blood. Patient muscle biopsies are characterized by atrophic myofibers with internalized nuclei, a hallmark shared with dystrophies and centronuclear myopathies^4^. Cardiac manifestations include arrythmia linked to late sodium current (L.B. Cronk, 2007), and atrio-ventricular conduction defects^5^. Even though no clear genotype-phenotype correlation exists, specific mutations have been linked to cardiac defects in addition to muscle symptoms, *in vitro* for F97C and S141R mutations^3^ and *in vivo* for P104L and T78M mutations^6,7^ . However, the underlying pathomechanisms of caveolinopathies is not fully understood, and to date there is no approved therapy for these diseases.

Caveolins are intramembrane proteins involved in the formation of caveolae, raft-like membrane invaginations that participates in various biological processes including endocytosis^8^, cell signaling^9,10^, mechanoprotection^11,12^ and lipid homeostasis^13,14^. Caveolin-3, which is exclusively expressed in striated muscles, was also shown to be located at nascent T-tubule^15,16^. T-tubules are membrane invaginations that facilitate rapid propagation of the action potential deep within the muscle fiber. Depolarization of the T-tubule membrane triggers activation of L-type calcium channels as dihydropyridine receptors (DHPRs; Ca_V_1.1) in skeletal muscle and Ca_V_1.2 in cardiac muscle, leading to the subsequent activation of ryanodine receptors—RyR1 in skeletal muscle and RyR2 in the heart— located on the sarcoplasmic reticulum. Once activated, these ryanodine receptors release calcium into the cytoplasm, initiating muscle contraction^17,18^. Several proteins participate in the biogenesis and maintenance of T-tubules in addition to caveolin-3, and include, non-exhaustively, amphiphysin 2 (BIN1), dynamin-2 (DNM2), dysferlin, and myotubularin (MTM1)^18,19^. Mutations in each of the corresponding genes leads to aberrant T-tubules structures in skeletal muscle and to a myopathic phenotype^18^. Cardiac T-tubules exhibit more complexity and form a network of transverse and longitudinal tubules with specific microdomains clustering ion channels. These microdomains are either formed by cardiac BIN1, caveolae or ankyrin B^20^. As in skeletal muscle, T-tubule impairment has a considerable impact on contractile function in heart and can notably be linked to heart failure and cardiomyopathy^21^.

A recent study focusing on T-tubule biogenesis during myofiber differentiation *in vitro* showed that caveolae composed of caveolin-3 form ring-like structures from which BIN1-positive tubules emerge^16^. This role in T-tubule biogenesis explains why T-tubules defects were described in a Cav-3 knock-out mouse model^22^. T-tubule defects can lead to impairment in the excitation-contraction coupling and to exercise intolerance in patients. Therefore, therapies aimed at restoring T-tubule function may offer a promising treatment strategy for caveolinopathies.

It was already demonstrated that human BIN1 overexpression, through transgenesis (*TgBIN1*), can rescue T-tubules and locomotor defects in mouse models for centronuclear myopathies^23,24^. Additionally, BIN1 is also able to spontaneously form membrane tubules when incubated alone with lipid bilayers^16,25^, suggesting that even in the absence of other T-tubule proteins such as caveolin-3, it might have the ability to create a network of T-tubules.

Here we hypothesize that BIN1 overexpression could improve T-tubule phenotype in a mouse model for caveolinopathy and, consequently, rescue muscle contraction defects.

Three murine model for caveolinopathies have been published: two knock-out models with a similar approach of exon 2 deletion causing mRNA decay^22,26^, and a knock-in model with the P104L mutation leading to sequestering of WT and mutant caveolin-3 in the Golgi^6^. The P104L model exhibit both cardiac and skeletal muscle phenotype including hypertrophic cardiomyopathy^6^, muscle atrophy and weakness^27^, with cellular dysregulation in mTORC1 signalling, cholesterol homeostasis^28^ and mitochondrial function^29^. Knock-out models were described with no alteration in growth and movement and no known effect on muscle force, but with increased muscle degeneration in the soleus and the diaphragm^22,26^, as well as progressive cardiomyopathy^30^, increased adiposity and insulin resistance^31^. Given that the *CAV3* mutations described in caveolinopathies exhibit a dominant-negative effect leading to a marked reduction or loss of caveolin 3 in patient muscles^4^, we choose the Cav-3 knockout mouse model established by Y. Hagiwara (2000) to investigate the loss-of-function mechanism common to all caveolinopathies. To overexpress BIN1 we selected the TgBIN1 mouse expressing the human *BIN1* gene, that was shown to completely replace murine BIN1 in a BIN1 muscle specific knockout^23,32^ .

Surprisingly, while BIN1 overexpression failed to restore locomotor function in skeletal muscle, it demonstrated significant therapeutic potential to rescue cardiac phenotypes.

## RESULTS

### Caveolin-3 loss induces muscle weakness

We first characterized further the *Cav-3* knockout mouse model. Marked *in situ* tibialis anterior (TA) weakness was observed at 3 months of age, in males and females, following sciatic nerve stimulations at both low and high frequencies (Fig. S1 A-D). This experiment reveals a muscle weakness in this mouse model and provides a strong muscular read-out to evaluate the impact of potential therapies.

*Cav-3*^-/-^ mice presented a distinct histological phenotype depending on the muscle studied. In the TA, few internalized nuclei were observed with no change in fiber diameter (Fig. S2 A), whereas a significant increase in nuclei internalization and percentage of small fibers was observed in the soleus at 3 months of age (Fig. S2 B). Overall, this mouse model reproduced the muscle weakness and the histopathological hallmarks described in patients.

### BIN1 overexpression in *Cav-3*^*-/-*^ does not rescue myopathic phenotype but restores myonuclei position

To assess if BIN1 overexpression could improve the motor and histological phenotypes of caveolinopathies, *Cav-3*^*+/-*^ mice were crossed with *BIN1* transgenic (*TgBIN1*) mice to obtain wild-type (WT), *Cav-3*^*-/*-^, *TgBIN1*, and *Cav-3*^*-/-*^ *TgBIN1* offsprings. BIN1 overexpression was confirmed with a 5-fold increase in protein level in mice with *BIN1* transgene in the TA (Fig. S3 A). The muscle-specific BIN1 isoform, that has a strong membrane tubulation property^33–35^, was found to be overexpressed by real time quantitative polymerase chain reaction in the TA and the heart (Fig. S3 B-C), and confirmed in the heart by western blot (Fig. S3 E). This isoform is typically found in human and rat heart^36^, while mouse heart mostly express the cardiac isoform (Fig. S3 D). To note, *TgBIN1* mice presented no locomotor phenotype compared to the WT (Fig. 1)^23^.

**Figure 1:**
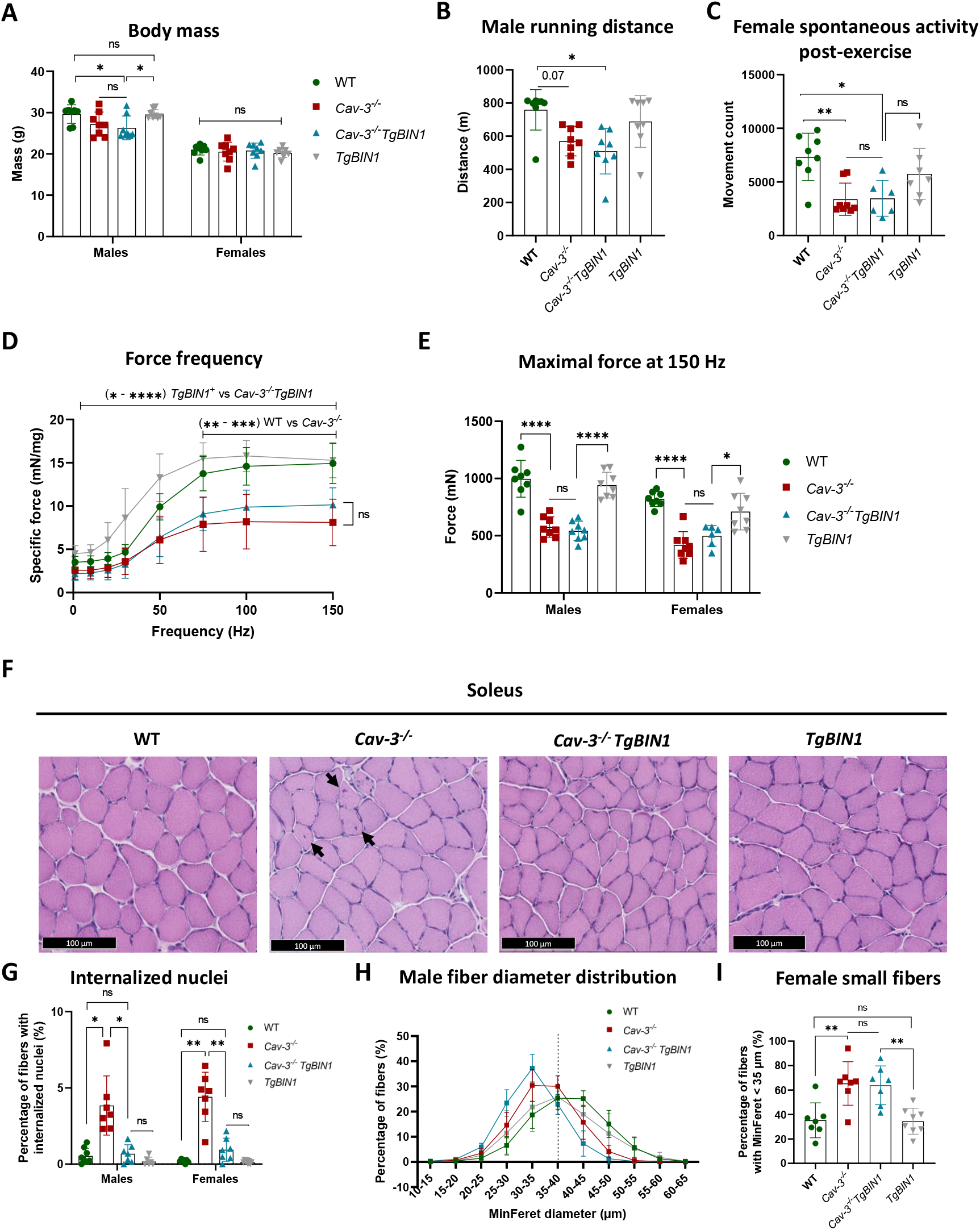
Overexpression of *BIN1* in Cav-3^-/-^ does not rescue skeletal muscle phenotype except nuclei positioning of 3 months old mice. (A) Body mass in males and females at 3 months of age. n=8. One-way ANOVA with Tukey’s multiple comparison (Males); Kruskal-Wallis with Dunn’s multiple comparison (Females). (B) Total distance run on a downhill treadmill by 3 months-old males. n=8; Kruskal-Wallis with Dunn’s multiple comparison test. (C) Total movement of female mice during 12 hours, monitored 8 hours after a downhill treadmill exercise. n=6-8; Kruskal-Wallis with Dunn’s multiple comparison test. (D) *In situ* force from male tibialis anterior (TA) muscle at increasing frequencies normalized to muscle mass. n=8; one-way ANOVA with Tukey’s multiple comparison or Kruskal-Wallis with Dunn’s multiple comparison at each frequency. (E) Absolute maximal TA force after sciatic nerve stimulation at 150 Hz. n=6-8; one-way ANOVA with Tukey’s multiple comparison. (F) Representative hematoxylin-eosin staining images of transversal soleus sections, with internalized nuclei indicated by black arrows. Scale bar = 100 μm. (G) Percentage of fibers containing at least one internalized nucleus in the soleus. n=6-8; Brown-Forsythe and Welch ANOVA with Dunnett’s T3 multiple comparison test. (H) Distribution of fiber MinFeret diameter in males soleus. n=6-7. (I) Percentage of fibers with a MinFeret diameter <35 μm in females soleus. n=7-8; One-way ANOVA with Tukey’s multiple comparison. *p<0.05; **p<0.01; ***p<0.001; ****p<0.0001.

*Cav-3*^*-/-*^ males presented a tendency toward decreased body mass and maximal running distance compared to WT, while females showed no difference in body mass but a significantly reduced spontaneous activity post exercise, based on actimetry measurements following a downhill treadmill protocol (Fig. 1 A-C). This phenotype reproduced the exercise intolerance that is the main sign of caveolinopathies. In both sexes, TA strength was strongly reduced (Fig. 1 D-E; Fig. S4C). *Cav-3*^*-/-*^*TgBIN1* showed decreased body mass in males compared to controls, reduced running abilities in males, and spontaneous activity in females, compared to WT. No difference in TA strength was observed between *Cav-3*^*-/-*^ and *Cav-3*^*-/-*^*TgBIN1* mice.

The impact of BIN1 overexpression on muscle histology was conducted on the soleus at 3 months of age. BIN1 overexpression efficiently corrected nuclei mispositioning in *Cav-3*^*-/-*^ mice (Fig. 1 F-G), but did not improve fiber atrophy in females and reinforced it in males (Fig. 1 H-I; Fig. S4E).

Overall, BIN1 overexpression was not sufficient to rescue locomotor and strength defects, as well as fiber atrophy in *Cav-3*^*-/-*^ mice. However, it efficiently corrected myonuclei positioning.

### BIN1 overexpression in *Cav-3*^*-/-*^ does not correct T-tubule ultrastructure in skeletal muscle

Our starting rationale was that BIN1 expression would promote T-tubule structure and rescue the motor defects. As we observed no improvement in muscle strength with BIN1 overexpression, we investigated whether BIN1 had an impact on T-tubule structure in the absence of caveolin-3. To do so, we examined T-tubule ultrastructure using electron microscopy in the TA, which exhibits measurable weakness, and in the soleus, which shows significant histopathological defects.

*Cav-3*^*-/-*^ T-tubules appeared swollen on certain sections of the tubules in both TA and soleus muscles (Fig. 2). In *Cav-3*^*-/-*^*TgBIN1* mice, we could also find swollen T-tubules, indicating that BIN1 overexpression did not rescue T-tubule defects.

**Figure 2:**
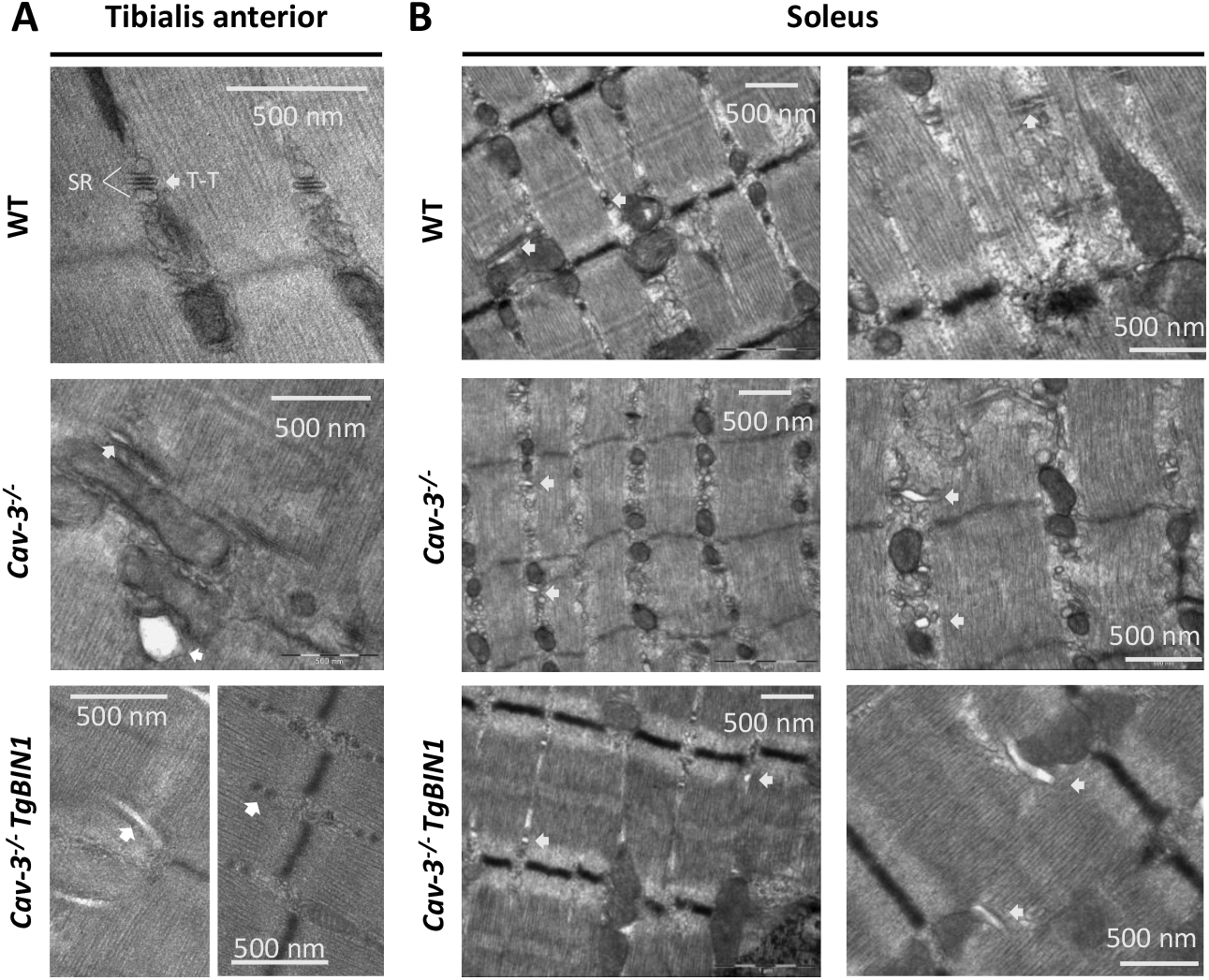
BIN1 overexpression in *Cav-3*^*-/-*^ mice does not improve T-tubule structure. (A-B) Electron microscopy on transversal muscle sections. Representative images of T-tubules (T-T) indicated by white arrows, and sarcoplasmic reticulum cisternae (SR) in the tibialis anterior (A) and soleus (B). Scale bar = 500 nm. (A) In the TA, both swollen (left) and normal T-tubules (right) are observed in *Cav-3*^*-/-*^ *TgBIN1*. n = 2-3.

In both soleus and TA muscles from *Cav-3*^*-/-*^ mice, we observed an abundance in degenerating mitochondria, with most of their inner content being electron-lucent with disrupted or lost cristae. These defects appeared less present in mice overexpressing BIN1, but not absent as in WT mice (Fig 3A & S5 A).

**Figure 3:**
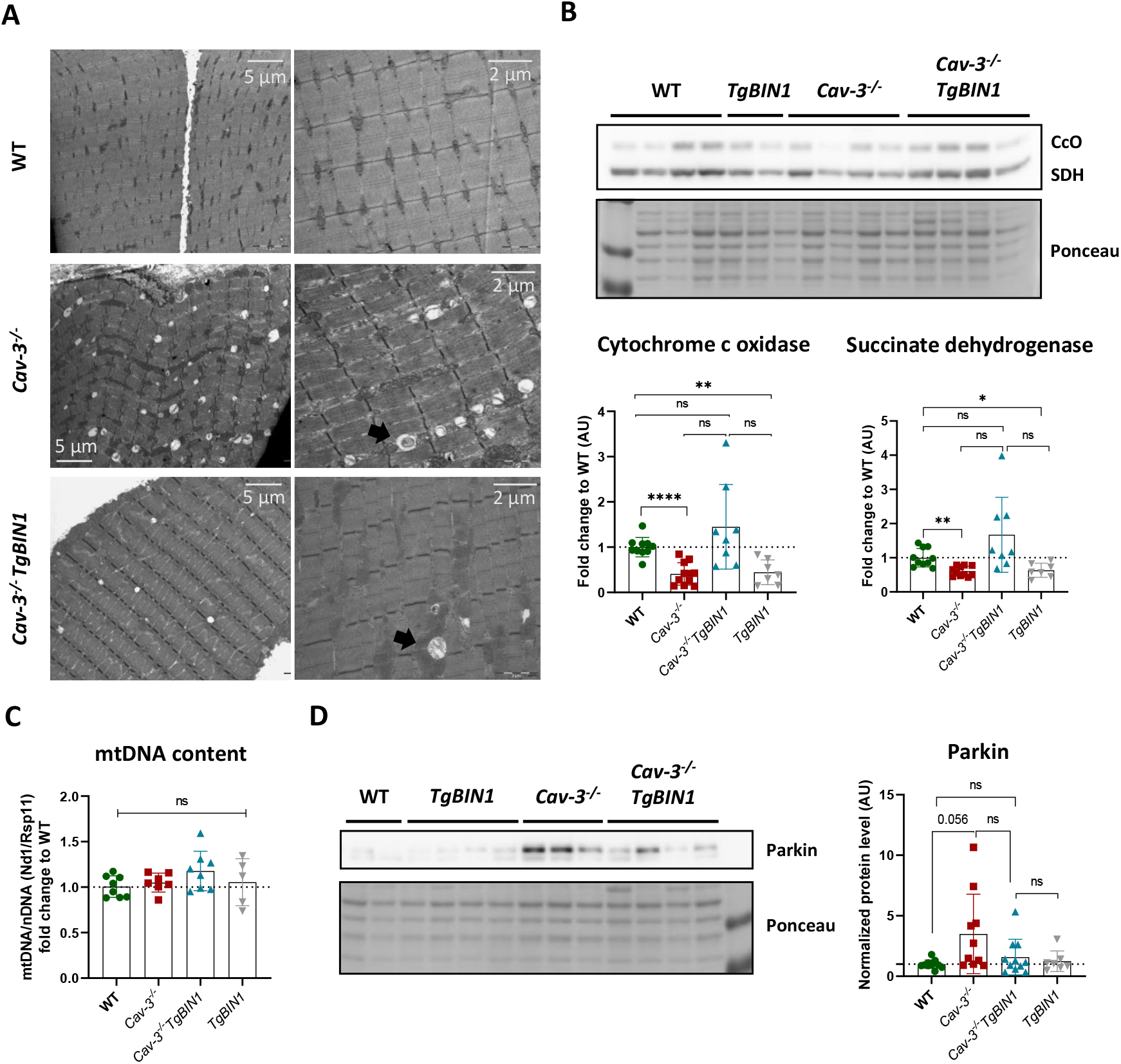
BIN1 overexpression in *Cav-3*^*-/-*^ mice has a beneficial effect on mitochondria homeostasis in the TA. (A) Electron microscopy on transversal muscle sections. Representative images of altered mitochondria indicated by a black arrow in TA muscle. Scale bar (left) = 5 μm. Scale bar (right) = 2 μm. n = 2-3. (B&D) Protein extracts from TA. Protein levels were normalized to Ponceau and represented as a fold change compared to WT. n= 7-11. Brown-Forsythe & Welch test with Dunnett’s T3 multiple comparison. (C) Total DNA extracts from TA. The ratio of *Nd1* mitochondrial gene on *Rsp11* nuclear gene estimates the mtDNA content. Data are represented as a fold change to WT. n = 5-8. One-way ANOVA with Tukey’s multiple comparison. *p<0.05; **p< 0.01; ****p<0.0001.

Although, no significant mitochondrial defects were observed with SDH staining in the soleus (Fig. S5 B), we detected a decrease in prohibitin protein level in *Cav-3*^*-/-*^ mice, that was improved with BIN1 overexpression (Fig. S5 C). In the TA, we observed decreased levels of succinate dehydrogenase (SDH) and cytochrome c oxidase (CcO) in *Cav-3*^*-/-*^ mice (Fig. 3B), consistent with the literature^29^, that could indicate an issue with the electron transport chain, as the mitochondria content is unchanged by caveolin-3 loss (Fig. 3C). Parkin protein level showed a tendency toward increase, hinting at a potential increase in mitophagy (Fig. 3E). BIN1 overexpression was able to improve CcO, SDH and Parkin levels in *Cav-3*^*-/-*^ mice.

Taken together, BIN1 overexpression did not correct T-tubules and motor defects in the skeletal muscle of *Cav-3*^*-/-*^ mice. However, it had a positive impact on nuclei positioning and mitochondria homeostasis.

### Caveolin-3 loss induces cardiac defects

Loss of function mutations in caveolin-3 are also associated with cardiac manifestations such as long QT syndrome or hypertrophic cardiomyopathy^2^, and a *Cav-3* knock-out mouse model was described with progressive cardiomyopathy^30^. We thus investigated in more detail the cardiac function of *Cav-3*^*-/-*^ mice and the impact of BIN1 expression. *Cav-3*^*-/-*^ male mice presented with reduced blood volume in the left ventricle during diastole (Fig. 4A), accompanied by reduced passive blood flow from the left atrium to the ventricle (Fig. 4B). These measures hinted at a diastolic dysfunction, such as a default in ventricle compliance, i.e. its capacity to expand upon blood arrival.

**Figure 4:**
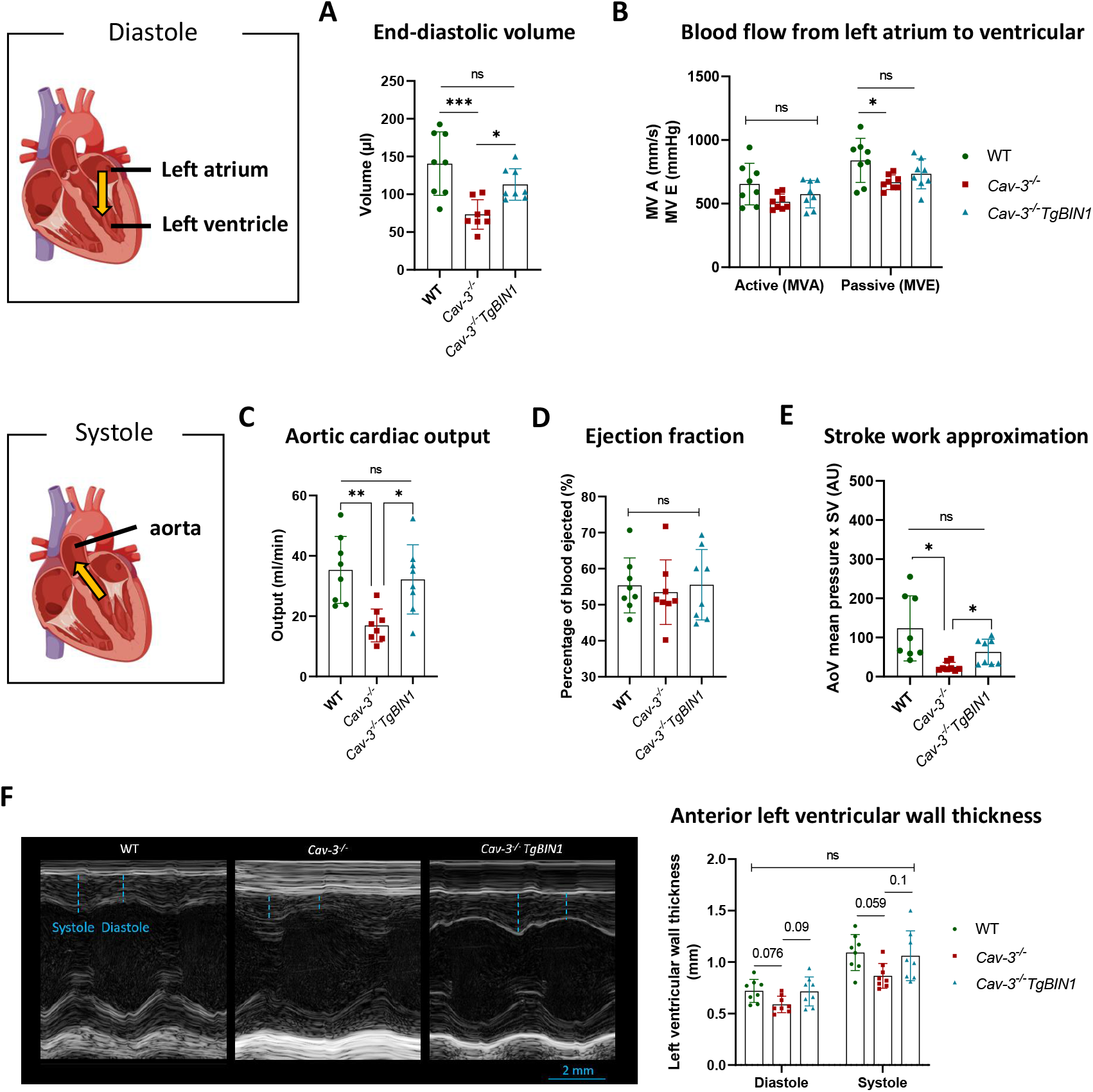
BIN1 overexpression in *Cav-3*^*-/-*^ mice rescues the cardiac function. (A-F) Echocardiography measurements in 3 months old males (n=8). (A) Volume of blood taken into the left ventricle at the end of the diastole. (B) Passive and active blood flow from the left atrium to the left ventricle. (C) Volume of blood ejected through the aorta per minute. (D) Percentage of the end-diastolic blood that is ejected during systole. (E) Approximation of the ability of the left ventricle to create pressure to eject the blood to the aorta. (F) Thickness of the anterior wall of the left ventricle. (A-D, F) one-way ANOVA with Tukey’s multiple comparison. (E) Brown-Forsythe and Welch ANOVA with Dunnett’s T3 multiple comparison. *p<0.05; **p< 0.01; ***p<0.001; ****p<0.0001.

Cardiac output, that depend on the blood available in the left ventricle, was decreased in *Cav-3*^*-/-*^ males, while the ejection fraction was unchanged, suggesting that the main cause for cardiac dysfunction could stem from diastolic issues (Fig. 4 C-D). However, future systolic dysfunction cannot be excluded, as both left ventricle contractile performance, showed by reduced stroke work, and structural integrity, indicated by a tendency toward wall thinning, appear affected in *Cav-3*^*-/-*^ males (Fig. 4 E-F).

Female *Cav-3*^*-/-*^ mice have a milder heart phenotype than males, with no apparent difference in diastolic function (Fig. S5 A-B), only a tendency toward decreased cardiac output and stroke work (Fig. S6 C-D), and no difference in left ventricular wall thickness (Fig. S6 E). Thus, we focused on males to study the impact of BIN1 modulation.

### BIN1 overexpression in *Cav-3*^*-/-*^ rescues cardiac function

Overexpression of BIN1 was previously shown to improve diastolic function in mice subjected to heart damaging stress^37^. Therefore, we investigated the impact of BIN1 overexpression in the heart of *Cav-3*^*-/-*^ mice using echocardiography. Here, BIN1 overexpression was able to rescue end-diastolic volume, cardiac output and stroke work (Fig. 4A-E). The cardiac performance index and the anterior left ventricular wall thickness were less impacted in the *Cav-3*^*-/-*^*TgBIN1* mice than in the *Cav-3*^*-/-*^ mice (Fig. 4F & S7 A). To characterize more in detail the aortic parameters, aorta diameter and aorta peak pressure were assessed and found decreased in *Cav-3*^*-/-*^ mice (Fig. S7 B-C). Noteworthy, these parameters were also improved following BIN1 overexpression. These data support a clear improvement of cardiac function in *Cav-3*^*-/-*^ mice overexpressing BIN1.

### BIN1 overexpression improves myofibril integrity in cardiomyocytes

To decipher how BIN1 overexpression could improve *Cav-3*^*-/-*^ cardiac phenotype, we investigated heart histology and ultrastructure. Reduced cardiac functions can be due to several factors including the accumulation of connective tissue, also called fibrosis. Fibrosis increases the stiffness of the tissues and could explain a decreased compliance of the left ventricle. *Cav-3*^*-/-*^ heart sections, stained for fibrosis with Masson’s trichrome, showed no difference compared to the WT controls (Fig. 5A). Similarly, hematoxylin-eosin staining did not reveal any defect in fiber organization in *Cav-3*^*-/-*^ heart sections (Fig. 5B).

**Figure 5:**
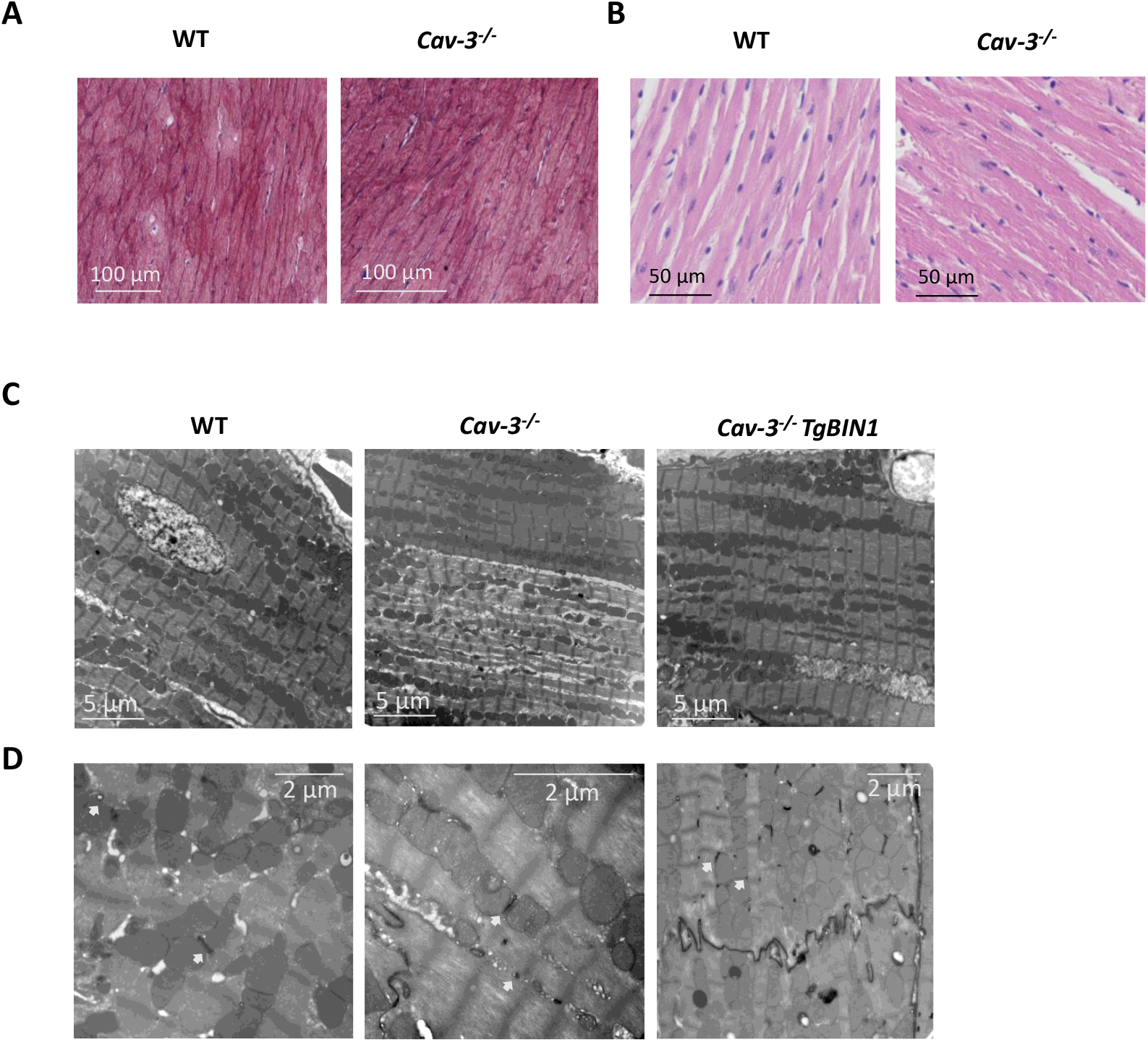
BIN1 overexpression in *Cav-3*^*-/-*^ mice improves sarcomere organization. (A) Representative images of Masson’s trichrome staining on transversal heart sections to assess fibrosis. Scale bar = 100 μm. (B) Representative images of hematoxylin-eosin staining showing cardiomyocyte morphology. Scale bar = 50 μm. (C-D) Electron microscopy on left ventricle sections. (C) Representative images of longitudinal sarcomere structure. Scale bar = 5 μm. (D) T-tubules visualized with potassium ferrocyanide staining (indicated by white arrows). Scale bar = 2 μm. n=3.

Electron microscopy of the left ventricle revealed that some *Cav-3*^*-/-*^ cardiomyocytes exhibited disconnected or thinned myofibrils, indicating compromised sarcomere integrity. These defects could not be found in WT and were rescued in *Cav-3*^*-/-*^*TgBIN1* ventricles (Fig. 5C). T-tubules structure marked by potassium ferrocyanide appeared unaffected by caveolin-3 loss with similar thin tubular shapes in the three genotypes (Fig. 5D).

Altogether, *Cav-3*^*-/-*^ heart present functional defects without observed anomalies in T-tubule structure. In this organ, overexpression of BIN1 was sufficient to improve myofibrils integrity, and to restore proper heart function.

### Loss of caveolin-3 has a differential impact on skeletal and cardiac muscles

To better understand why BIN1 overexpression affected cardiac and skeletal muscles differently, and to explain the differential impact of caveolin-3 loss on T-tubules structure between both tissues, we compared the expression of genes coding for proteins implicated in caveolin-related functions and in excitation-contraction coupling in the heart and tibialis anterior (TA) using RT-qPCR.

The tibialis anterior appeared globally more impacted by caveolin-3 loss than the heart (Fig. 6A), with caveolae component such as Cavin-1 and -4 being upregulated at the gene expression level (Fig. 6B). As cavin-4 protein level is within normal range in these tissues (Fig. 6F), it suggests that the increased mRNA expression is a compensatory mechanism for altered cavin-4 function or level due to lack of caveolin-3. In addition, the ryanodine receptor expression was decreased in TA muscle while the L-type calcium channel expression (*Cacna1s*) was increased, suggesting a stronger dysregulation of excitation-contraction coupling in skeletal muscle (Fig. 6C). Moreover, expression of the acetylcholine receptor subunits was abnormal in TA muscle, potentially impacting contraction (Fig. 6E). The observed skeletal muscle phenotype was linked with a significant increase in atrogenes expression such as *MuRF1*, a tendency toward increase for *Atrogin1* (Fig. S8) and an increase in *Gadd45a* expression that is generally associated with muscle weakness, atrophy and mitochondria defects^38^ (Fig. 6D).

**Figure 6:**
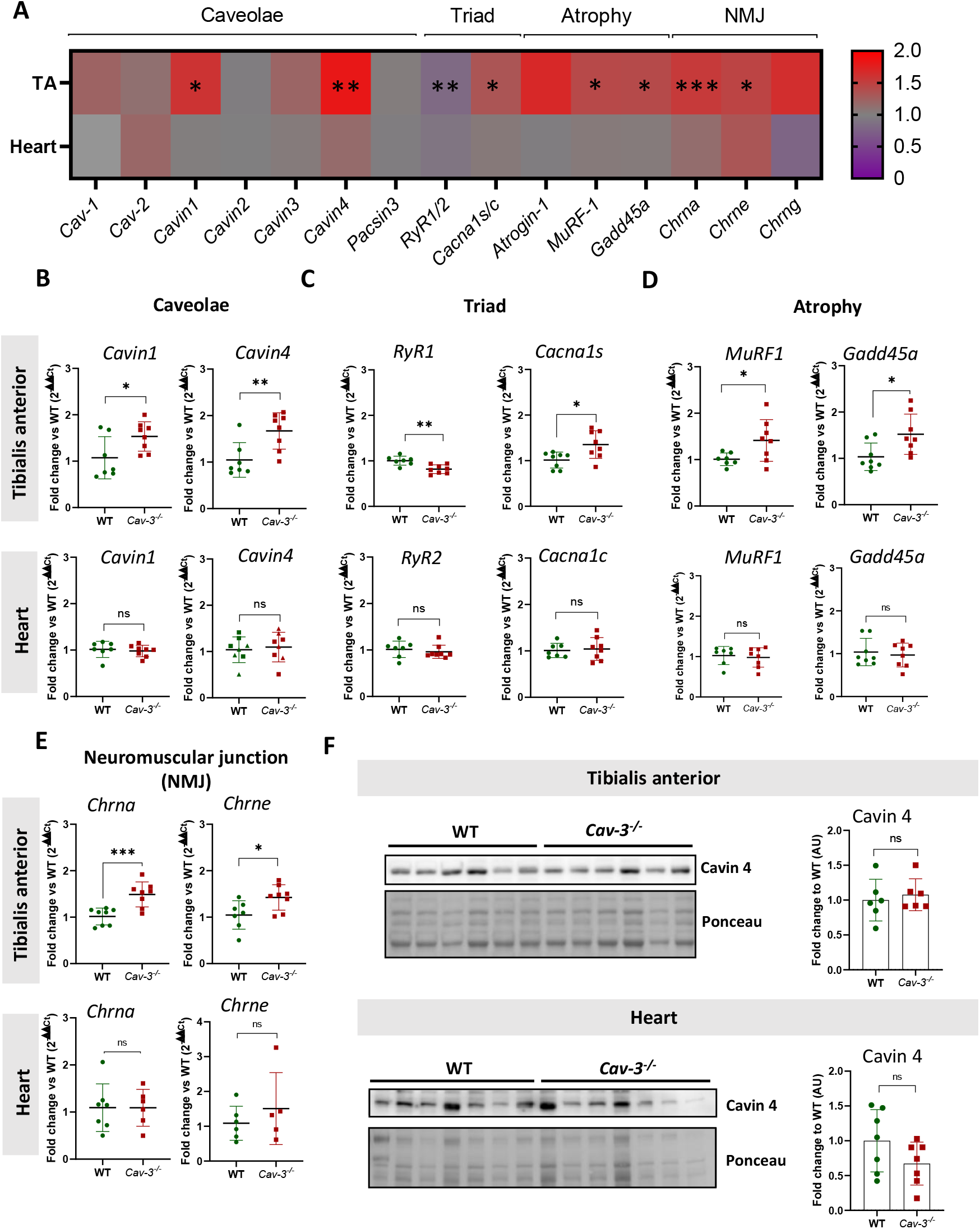
Loss of caveolin-3 has a differential impact on skeletal and cardiac muscle. (A) Heatmap representation of relative mRNA expression levels for genes involved in caveolae structure, excitation–contraction coupling, and muscle atrophy pathways, measured in the tibialis anterior (TA) and heart of WT and *Cav-3*^*-*^*/*^*-*^ mice by RT-qPCR. Quantification of transcript levels for caveolae-related proteins as cavins (*Cavin1–4*) (B), excitation-contraction coupling regulators (*RyR1/2, Cacna1s/c*) (C), muscle atrophy-related genes (*MuRF1, Gadd45a*) (D), and neuromuscular transmission (*Chrna/e*) (E) in TA muscle and heart. Data are presented as fold change relative to WT. n=5-8; Unpaired Student’s t-test or Mann-Whitney; \**p*<0.05; ** p<0.01; ***p<0.001.

In conclusion, these molecular alterations underlie the structural and contractile defects observed in skeletal muscle. Moreover, the more pronounced impairment in skeletal muscle compared to cardiac muscle may explain why BIN1 overexpression is more effective in rescuing cardiac, rather than skeletal phenotypes.

## DISCUSSION

In this study, we evaluated the therapeutic potential of BIN1 overexpression in a mouse model of caveolinopathy. We found that *Cav-3*^*-/-*^ mice display both skeletal muscle weakness and cardiac dysfunction, with altered T-tubule morphology in muscle and preserved T-tubules but functional defects in the heart. While BIN1 overexpression failed to rescue skeletal muscle force, T-tubule architecture, or fiber atrophy, it successfully restored nuclei positioning and improved mitochondrial markers in myofibers. Importantly, BIN1 expression markedly improved cardiac function and sarcomeric organization. The differential rescue between heart and muscle shed light on different impact of caveolin-3 loss in the two striated muscle types.

### Skeletal and cardiac muscle duality

Caveolinopathies are complex disorders affecting both cardiac and skeletal muscle, and caveolin-3 plays key roles in the membrane architecture sustaining excitation-contraction coupling, and in mechano-signaling. Prior studies demonstrated that BIN1 can restore T-tubule structure and function in models of centronuclear myopathies^23,24,39^. Of note the histopathological hallmarks of caveolinopathies and centronuclear myopathies are very similar, with atrophic myofibers with internal nuclei and lack of obvious dystrophic features in patients^4^. Based on this, we hypothesized that BIN1 overexpression might also correct defects in caveolinopathies, particularly by targeting the T-tubule disorganization observed in *Cav-3*^*-/-*^ muscle fibers. Our findings highlight a striking divergence in the response to BIN1 overexpression between skeletal and cardiac muscle. This duality may stem from two non-mutually exclusive mechanisms.

First, distinct pathomechanisms may underlie skeletal versus cardiac defects in caveolinopathies. In skeletal muscle, the pathology appears primarily driven by T-tubule disorganization, in addition to mechano-transduction investigated in other studies^11^. This is supported by the swollen and disorganized T-tubules observed in *Cav-3*^*-/-*^ muscle and the lack of force restoration despite BIN1 overexpression. Comparable T-tubule swelling has been reported in other T-tubule-related disorders, such as *Mtm1*-or *Bin1*-deficient mice^40^ and patients with centronuclear myopathy^41^, and *MG29* knockout mice^18,42^ . Gene expression analyses also revealed a stronger deregulation of genes related to excitation-contraction coupling in the tibialis anterior compared to the heart. Alternatively, muscle weakness may be linked to myofiber atrophy, a hallmark present in the *Cav-3*^*-/-*^ muscle and in patients, not rescued by BIN1 overexpression. Myostatin or TGFb1 receptor inhibition was able to improve muscle atrophy in Cav-3^-/-^ mice^27,43^, while BIN1 overexpression does not induce muscle hypertrophy in mice^23^. Conversely, the cardiac phenotype in *Cav-3*^*-/-*^ mice, marked by reduced diastolic filling, occurred without major T-tubule defects, suggesting that ion channel mislocalization or dysregulation, possibly involving Kir2.1 or L-type calcium channels, may contribute more strongly in the heart. It is possible that reduced end-diastolic volume is due to impaired ventricular repolarization, the latter being recently linked to caveolin-3^44^.

A second hypothesis is that the BIN1 rescue requires tissue-specific co-factors. Skeletal muscle expresses high levels of caveolin-3 and cavin-4, both of which are necessary for triad formation. In contrast, cardiac muscle co-expresses caveolins-1 and -2, which may functionally compensate for the loss of caveolin-3. Indeed, a recent study suggests that the *CAV-3* T78K mutation would be incompatible with life without caveolin-1 presence^45^ . Additionally, the double Cav-1/Cav-3 knockout mouse develops a more severe cardiac phenotype^46^. The presence of these caveolin paralogs may permit BIN1 to stabilize membrane-cytoskeletal structures in the heart, even in the absence of caveolin-3. Cavin 4 also represents an important co-factor for BIN1 in skeletal muscle. Cavin-4 has been shown to bind and cooperate with BIN1 at T-tubules^47^. We found that cavin-4 expression is dysregulated specifically in skeletal muscle of *Cav3*^*-/-*^ mice, but not in heart. Previous studies showed a loss of caveolae in Cav-3 knock-out mouse model^26^. Losing its scaffold could impact cavin 4 stability or function, and would explain, why the increase in gene expression is not linked to cavin 4 protein increase in *Cav-3*^*-/-*^ mice. Consequently, cavin-4 defects may prevent BIN1 from properly executing its membrane-shaping role at the T-tubules.

Together, these findings emphasize that the pathomechanisms underlying caveolinopathy are tissue-specific and that effective rescue may depend on the availability of cofactors, structural scaffolds, and distinct cellular stress responses.

### Translational implication for BIN1 gene therapy

Our work reinforces the growing body of evidence supporting BIN1 as a therapeutic gene for cardiac disease. Previous studies have demonstrated the benefits of AAV9-mediated BIN1 delivery in models of heart failure, stress-induced cardiomyopathy, and diabetic heart disease^37,48–50^ . Here, we extend these findings to a genetic cardiomyopathy caused by caveolin-3 deficiency. The cardiac rescue observed in our *Cav-3*^*-/-*^ mice further suggests that muscle-specific BIN1 isoforms can confer structural and functional benefits in the heart, possibly by preserving sarcomeric integrity.

While BIN1 alone can tubulate membranes in vitro and in some disease models, its function appears insufficient in the absence of structural partners such as caveolin-3, and potentially cavin-4. This suggests the need for combination strategies, such as co-delivery of scaffolding factors or targeting upstream regulators of triad assembly, for treating skeletal muscle manifestations in caveolinopathies.

### Conclusion

Overall, this study demonstrates that BIN1 overexpression selectively rescues cardiac but not skeletal muscle defects in a caveolinopathy mouse model. These findings highlight the tissue-specific consequences of caveolin-3 loss and reinforce the importance of BIN1 as a therapeutic candidate for cardiac disease, while highlighting the importance of exploring new approaches in skeletal muscle.

## MATERIAL AND METHODS

### Mouse models and ethical approval

This study was conducted in accordance with the French legislation under the APAFIS project number: 2022111011252590. The Cav-3 knockout or Cav-3^tm1Ncnp^ mouse model was developed by Toshikuni Sasaoka, published in Y. Hagiwara in 2000 and supplied by RIKEN on the C57BL/6J100% background. TgBIN1 mice were created through the insertion of human BAC (no. RP11-437K23; Grch37 chromosome 2: 127761089-127941604) encompassing the full *BIN1* gene and were previously described in V.M. Lionello, 2019.

### Locomotor phenotyping

The tests were performed by the same experimenter, who was blinded to the genotype.

Locomotor tests were performed at 3 months of age and consisted of two sequential protocols. First, locomotor ability during forced exercise was assessed using a treadmill apparatus from Bioseb with a -15° inclination for a 3 phase - 50 mins running test, maximum. Exercise started with an endurance phase of 30 mins at 24 cm/s, followed by intense exercise with increasing speed over shorter period of time: 33 cm/s – 5 mins; 36 cm/s – 5 mins; 40 cm/s – 1 min; 44 cm/s – 1 min, and finally a sustained exercise at 24 cm/s for 7 mins with a 1 min transition at 30 cm/s between intense and sustained phases.

Each running track has an electrical grid delivering mild electric shock (0.2 mA) to encourage running by negative stimuli. Habituation was done earlier in the day for 5 mins at 10 cm/s with -15° inclination.

During the night following the treadmill, spontaneous activity was measured by an actimeter (Imetronic). Each mouse is placed in an individual cage with food and water ad libitum. Rearing and spontaneous movement are recorded over 12 hours during their active phase. Mice are placed one hour before the start of the experiment to let them settle in, but also to limit the time spent in isolation.

### *In situ* muscle force

The tests were performed by the same experimenter. As this test precedes organ collection into annotated tubes, awareness of the genotype cannot be excluded.

Tibialis anterior force measurements were done on the Complete 1300A Mouse Test System from Aurora Scientific. The experiment was conducted under general anesthesia with a mix of (1) Domitor/Fentanyl (2/0.28 mg/kg), (2) diazepam (8 mg/kg) and (3) Fentanyl (0.28 mg/kg) 15 mins after the first two injections, to achieve narcosis, sedation, analgesia and myorelaxation. Cutaneous and muscular plan are open near the hip to expose the sciatic nerve. The lower tendon of the tibialis anterior is sectioned and attached to a transducer using nylon thread. The tibialis anterior is partly detached (3/4) and the knee and paw are fixed. Sciatic nerve is stimulated at low (2 Hz) to high (150 Hz) frequencies. Specific force is calculated by normalizing to the tibialis anterior mass.

### Cardiac phenotyping

The tests were performed and analysed by the same experimenter, who was blinded to the genotype.

Echocardiography was performed at 7 weeks of age. Anesthesia was induced with 3-4 % isoflurane and maintained at 1-2%. Body temperature was maintained using a heat pad and heat lamp and monitored with a rectal thermometer. Tests were performed as previously described^51^ using Chart 4.2.3 software, an electrocardiograph ISO DAM8 amplifier (World Precision Instruments, USA) and analogic-numeric conversion box (ITF16A/D converter, EMKA technologies, USA) for electrocardiography, and a VisualSonics Vevo 2100 Imaging System (Toronto, Canada) with a MS400 probe (30-MHz) for echocardiography.

### Tissue collection and histology

Tibialis anterior and soleus were dissected and frozen in liquid-nitrogen cold isopentane. 8 μm transversal cryosections were obtained using the Leica CM3050 S cryostat and stained, by the ICS histology platform, with HE, SDH and Masson’s trichrome. Slides images were acquired on the Axioscan 7 from Zeiss and the Cellpose algorithm was used to segment individual fibers. Quantifications were done on Fiji manually using CellCounter. One transversal section was quantified for each animal.

### Electron microscopy

Directly after dissection, muscles and left ventricles were cut in cacodylate buffer (0.1 M, pH 7.4) into 1x1 mm fragments, and then transferred into 2.5% glutaraldehyde and 2.5% paraformaldehyde cacodylate buffer for fixation at 4°C overnight.

Skeletal muscle: Fixed samples were rinsed and kept in 1% osmium tetroxide reduced with 1.5% potassium hexacyanoferrate (III) [K_3_Fe(CN)_6_] in 0.1 M cacodylate buffer for 1 h at 4 °C. Samples were dehydrated through a graded ethanol series (50%, 70%, 90%, and 100%) and propylene oxide, each for 30 min, then oriented and embedded in Epon 812 resin. Semithin sections (2 μm) and ultrathin sections (70 nm) were cut using a Leica Ultracut UCT ultramicrotome. Sections were contrasted with uranyl acetate and lead citrate, and examined at 70 kV using either a Morgagni 268D electron microscope (FEI Electron Optics) equipped with a Mega View III camera (Soft Imaging System).

Heart: After rinsing in distilled water, samples were post-fixed for 2 h at 4 °C in 2% osmium tetroxide reduced with 1.5% potassium hexacyanoferrate (III) [K_3_Fe(CN)_6_] in distilled water. Following extensive water rinses, tissues were post-stained with 1% uranyl acetate for 2 h at room temperature and rinsed again in water. Samples were dehydrated in a graded ethanol series (25%, 50%, 70%; 30 min each; 2x 90%, and 3 × 100%; 1 h each) and and left overnight in ethanol/EPON resin 4:1 at RT with a gentle shake. Then samples were infiltrated with a graded ethanol/EPON resin series (ethanol:EPON = 2:1, 1:1, 1:2, and finally 2 × 100% resin). Polymerization was carried out at 60 °C for 48 h. Ultrathin sections (70 nm) were mounted on grids and examined using a Hitachi H7500 transmission electron microscope operating at 80 kV, equipped with an AMT Hamamatsu digital camera.

Acquisition was performed by the same experimenter for each organ. Both were blinded to the genotype.

### RNA extraction and RT-qPCR

Frozen tissues were lysed mechanically with the Precellys® Evolution Touch system from Bertin Technologies (2x 15 s 5500 rpm pulses) in TRI Reagent (#TR118, Molecular Research Center) . Total RNA was recovered using chloroform, precipitated with isopropanol and washed with ethanol 75% before resuspension in DEPC-treated water. Complementary DNA (cDNA) was synthesized using SuperScript IV Reverse Transcriptase (#18090010, Invitrogen) and quantitative real-time PCR was carried out using SYBR Green Master Mix I (#04887352001, Roche Diagnostics) on a Lightcycler 480 system (Roche Diagnostics). Reactions included biological and technical replicates (3-4 replicates). Primers can be found in the Supplementary Table 1.

### Protein extraction and western blotting

Total protein was extracted in RIPA buffer (150 mM NaCl, 50 mM Tris (pH 8), 0.5% sodium deoxycholate, 1% NP-40, and 0.1% SDS) supplemented with 1 mM PMSF, 1 mM sodium orthovanadate, 5 mM sodium fluoride, and 1X protease inhibitor cocktail and 0.2 mM DTT. Tissue samples were lysed mechanically using a Precellys® Evolution Touch tissue homogenizer from Bertin Technologies (2x 20 s at 6000 rpm pulses) and supernatant was kept after a 5 mins 10 000 g centrifugation. Protein concentrations were determined using the DC Protein Assay Kit (#5000116, BioRad). Protein were reduced in Pierce Lane Marker buffer (Thermofischer #39000) and heated at 95°C for 5 mins before loading on the gel (or 50°C for OxPhos cocktail). Polyacrylamide gels were prepared using standard protocol. 12 μg of proteins were loaded per lane. Transfer on nitrocellulose membrane was performed using the Trans-Blot Turbo RTA Mini Nitrocellulose Transfer Kit (#170-4270, BioRad) for 10 mins. Membranes were stained with Ponceau for total protein quantification. Following 1 h blocking in 5% nonfat dry milk in 0.1% Tween20-TBS (TBST), membranes were incubated overnight at 4°C with primary antibody diluted in 5% milk-TBST. Membranes were rinsed 3 times 5 mins in TBST and were incubated for 1 h in secondary antibody in 5% milk-TBST. Images were acquired on an Amersham Imager 600 (GE Healthcare Life Sciences) after a 2 mins incubation in chemiluminescent reagent (Immobilon HRP substrate Millipore #WBKLS0100). Quantification was performed on Fiji and data were normalized to the Ponceau and the mean value of the control group to obtain fold change values. Primary antibodies are listed in Supplementary Table 1. Uncropped images are available in Supplementary Material 1.

### Statistical analysis

Statistical analyses and graph generation were conducted in GraphPad Prism (v10.0.2). Data normality was evaluated using the Shapiro–Wilk or the D’Agostino–Pearson test (n≥8). For normally distributed datasets with homogeneous variances, one-way ANOVA was performed. When variances were unequal, Brown–Forsythe and Welch ANOVA were used. Log-normally distributed data were log-transformed prior to ANOVA. Non-normally distributed data were analyzed using the Kruskal–Wallis test. For each ANOVA, multiple comparison was performed with the mean of each group being compared to the mean of every other group. Significance was defined as p < 0.05. Graphs present individual data points as well as mean ± SD. Each value is available along with statistical methods in Supplementary Material 2.

### Inclusion and exclusion

Mouse genotype was verified again before histological and molecular analysis. In the case of prior genotyping error, animals were moved, when possible, to the correct group for the *in vivo* analysis or excluded. In the case of freezing defects observed in histology, animals were removed from this specific analysis. Each exclusion is specified in Supplementary Material 2.

## Data availability

All source data supporting the findings of this study are provided with the article.

## Acknowledgements

We thank Ghina Bou About for her cardiac expertise, and acknowledge the support of the cardiology, histology, animal care and electron microscopy platforms at the Institut de Génétique et de Biologie Moléculaire et Cellulaire (IGBMC).

